# Protection of rhesus macaques against vaginal SHIV challenges by VRC01 and an anti-α_4_β_7_ antibody

**DOI:** 10.1101/365551

**Authors:** Giulia Calenda, Ines Frank, Géraldine Arrode-Brusés, Amarendra Pegu, Keyun Wang, James Arthos, Claudia Cicala, Brooke Grasperge, James L. Blanchard, Stephanie Maldonado, Kevin Roberts, Agegnehu Gettie, Anthony S. Fauci, John R. Mascola, Elena Martinelli

## Abstract

VRC01 protects macaques from vaginal SHIV infection after a single high-dose challenge. Infusion of a simianized anti-α_4_β_7_ mAb (Rh-α_4_β_7_) just prior to, and during repeated vaginal exposures to SIVmac251 partially protected macaques from vaginal SIV infection and rescued CD4^+^ T cells. To investigate the impact of combining VRC01 and Rh-α_4_β_7_ on SHIV infection, 3 groups of macaques were treated with a suboptimal dosing of VRC01 alone or in combination with Rh-α_4_β_7_ or with control antibodies prior to the initiation of weekly vaginal exposures to a high dose (1000TCID_50_) of SHIV_AD8-EO._ The combination Rh-α_4_β_7_-VRC01 significantly delayed SHIV_AD8-EO_ vaginal infection. Following infection, VRC01-Rh-α_4_β_7_-treated macaques maintained higher CD4^+^ T cell counts and exhibited lower rectal SIV-DNA loads compared to the controls. Interestingly, VRC01-Rh-α_4_β_7_-treated macaques had less IL-17 producing cells in the blood and the gut during the acute phase of infection. Moreover, higher T cell responses to the V2-loop of the SHIV_AD8-EO_ envelope in the VRC01-Rh-α_4_β_7_ group inversely correlated with set point viremia. The combination of suboptimal amounts of VRC01 and Rh-α_4_β_7_ delayed infection, altered anti-viral immune responses and minimized CD4^+^ T cell loss. Further exploration of the effect of combining bNAbs with Rh-α_4_β_7_ on SIV/HIV infection and anti-viral immune responses is warranted and may lead to novel preventive and therapeutic strategies.

**Short summary:** A combination of VRC01 and Rh-α_4_β_7_ significantly delayed SHIV acquisition, protected CD4 counts, decreased gut viral load and modified the immune response to the virus.

## INTRODUCTION

Integrin α_4_β_7_ (α_4_β_7_) is expressed at high levels by CD4^+^ T cells trafficking to the gut associated lymphoid tissues (GALT) ^1-3^, a critical site for HIV replication and dissemination after transmission ^4-7^. α_4_β_7_^high^ CD4^+^ T cells are highly susceptible to HIV infection and are preferentially depleted during acute HIV-1 and SIV infection ^8-10^. Higher frequencies of α_4_β_7_^high^ CD4^+^ T cells have been correlated with increased susceptibility to HIV-1 infection in humans and SIV infection in macaques and with disease progression in both humans and macaques ^11,12^. The higher risk of HIV-1 acquisition due to prevalent HSV-2 infection has also been associated with increased levels of α_4_β_7_ expression ^13-15^. Targeting α_4_β_7_ with a simianized anti-α_4_β_7_ monoclonal antibody (Rh-α_4_β_7_; mAb) prior to and during a repeated low-dose vaginal challenge (RLDC) study in rhesus macaques prevented SIV acquisition in half of the animals and delayed disease progression in those animals that did become infected ^16^. Reportedly, simultaneous treatment with Rh-α_4_β_7_ and cART led to sustained viral control after cessation of all forms of therapy in at least one model of SIV infection ^17^. The mechanism(s) underlying the anti-HIV-1 activity of the Rh-α_4_β_7_ mAb are poorly understood. Rh-α_4_β_7_ does not block viral entry into CD4^+^ T cells and has weak anti-HIV-1 activity *in vitro* ^8,18,19^. We have recently shown that signaling through α_4_β_7_ can promote HIV replication^20^ and, in this regard, we previously demonstrated that Rh-α_4_β_7_ blocks α_4_β_7_ from adopting an active conformation that is critical for this signaling ^21^.

In addition, we determined that Rh-α_4_β_7_ selectively alters trafficking of CCR6^+^ CD4^+^ T cells to mucosal tissues ^22^ and impacts the antibody response to SIV infection when given in combination with cART ^17^. Thus, interference with both immune cell trafficking and α_4_β_7_-driven viral amplification may, at least in part, explain the decrease in gut tissue SIV loads when Rh-α_4_β_7_ is administered prior to, and throughout the acute phase of infection^23^.

Passive transfer of a number of broadly neutralizing antibodies (bNAbs) targeting HIV-1 envelope (Env) has been shown to protect rhesus macaques against a single high-dose inoculation with simian-human immunodeficiency virus (SHIV) ^24-27^ and this strategy is currently being evaluated to prevent HIV-1 acquisition in humans ^28^. In particular, VRC01, a bNAb against the CD4 binding site (CD4bs) on the HIV envelope ^29,30^, is the first bNAb to be investigated clinically for the prevention of HIV infection in adult men and women (AMP trial; NCT02716675 and NCT02568215). Moreover, VRC01 is being tested for safety in HIV-exposed infants (NCT02256631) as a potential agent to prevent mother-to-child transmission (MTCT) of HIV-1. In preclinical studies, VRC01 protected monkeys against single high-dose vaginal and rectal SHIV challenge ^27^ and its protective activity against repeated low-dose rectal challenges decreases after several weekly challenges ^31^. In this regard, bNAb protection against repeated rectal challenges was shown to be dependent on the potency and half-life of bNAbs ^31^. A mutation in the Fc domain of the antibody, which was shown to increase VRC01 half-life in both plasma and tissues, increased ^32^ and prolonged ^31^ its protective activity. Several other strategies to improve the pharmacokinetics and function of bNAbs ^28^ as well as the use of combinations of bNAbs or bi- and trispecific antibody-based molecules ^33-35^ are being tested with the ultimate goal of generating new prevention and therapeutic options against HIV-1 infection.

In the present study, we investigated the combination of VRC01 and Rh-α_4_β_7_ in a repeated vaginal challenges model using the tier 2 SHIV_AD8-EO_ ^36^. This challenge virus was chosen for its multiple properties typical of pathogenic HIV-1 isolates ^37^, allowing to explore the impact of the VRC01-Rh-α_4_β_7_ combination on SHIV_AD8-EO_ infection and anti-viral immune responses during the acute and early chronic phase of infection. In order to detect an effect of this combination over the sterilizing protective effect of VRC01, we chose a repeated high-dose model of infection and treatment with suboptimal amounts of both antibodies. The VRC01-Rh-α_4_β_7_ combination significantly delayed SHIV_AD8-EO_ acquisition, protected blood CD4^+^ T cells and altered anti-viral immune responses.

## RESULTS

### The VRC01-Rh-α_4_β_7_ combination significantly delays SHIV_AD8-EO_ vaginal infection

VRC01 has been shown to provide sterilizing protection against high-dose vaginal challenge with SHIV_SF162P3_ ^27,38^. In order to study the VRC01-Rh-α_4_β_7_ combination in a setting of suboptimal VRC01 protection we employed an inoculum 100 fold higher and half the dose of VRC01 (10mg/kg) that resulted in delayed SHIV_AD8-EO_ acquisition for a median of 8 weeks in a repeated rectal low-dose challenge model ^31^. A VRC01-alone group was used to monitor baseline VRC01 protection.

A total of 27 animals were infused with VRC01 (10mg/kg) and Rh-α_4_β_7_ (25mg/kg; n=9) or with 10mg/kg of VRC01-alone (n=9) or with control human and rhesus IgGs (n=9), 3 days before weekly vaginal challenges with a high-dose inoculum of SHIV_AD8-EO_ (1000TCID_50_) until all animals became infected (Fig 1A). Rh-α_4_β_7_ infusions were repeated every 3 weeks for a total of 6 infusions. A Rh-α_4_β_7_-alone group was not included because Rh-α_4_β_7_ does not protect from high-dose challenge^39^ and the levels of VRC01 rapidly decrease^27^. Thus, the impact of Rh-α_4_β_7_ on SHIV-AD8 infection can be inferred from comparison with the VRC01-only and control groups.

**Figure 1.**
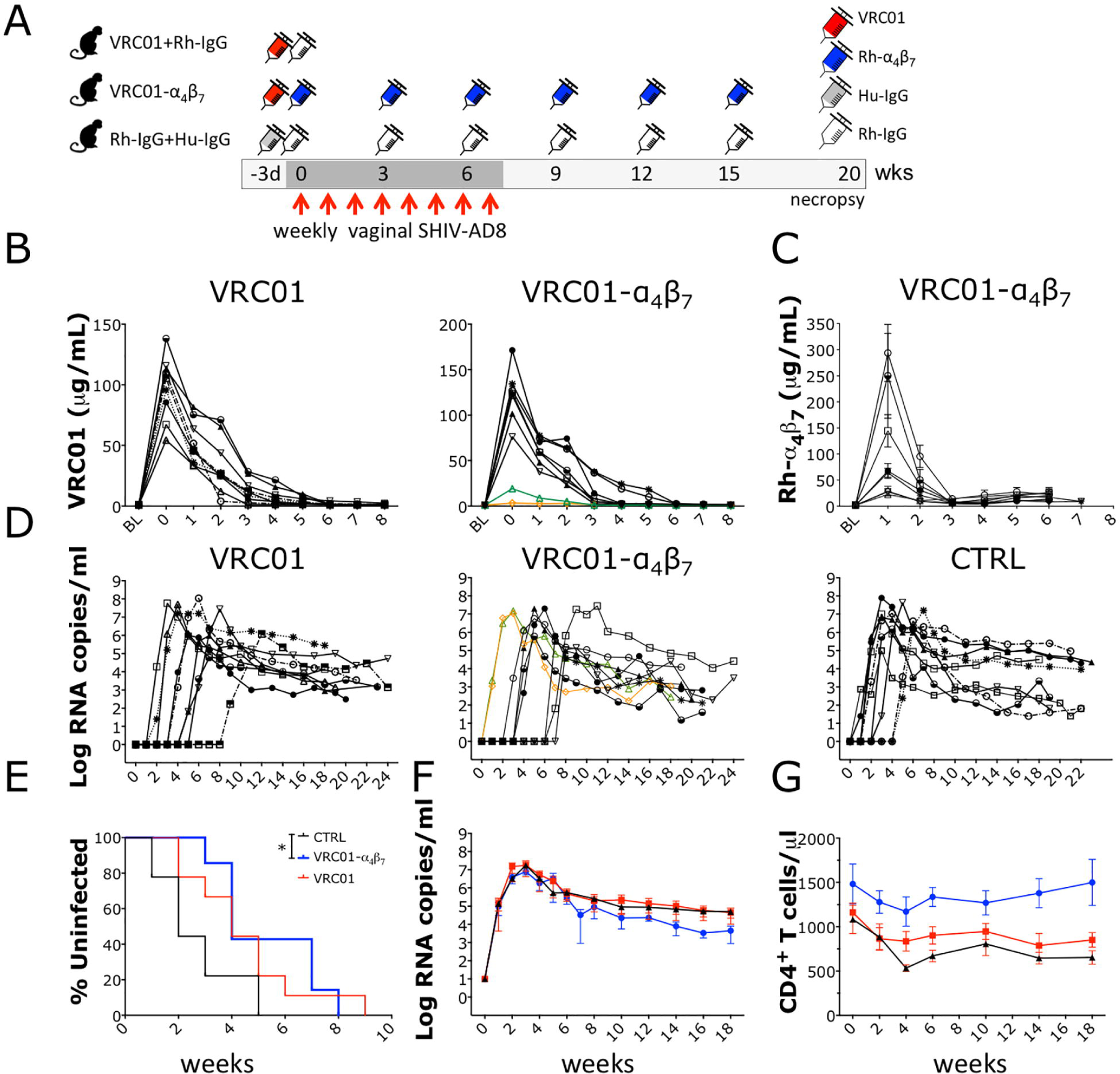
The Rh-α_4_β_7_-VRC01 combination significantly delays SHIV_AD8-EO_ acquisition. (A) Schematic of the study procedures (B-C) The concentrations of VRC01 (B) and Rh-α_4_β_7_ (C) were measured in plasma for the first 8 weeks after initiation of treatment. VRC01 levels were below the protective concentration in GH63 and KT57 (orange and green in B, respectively). (D) Copies of viral RNA copies in plasma are shown from the time of the first challenge (week 0). (E) Kaplan-Meier curves generated with time to first viral detection in plasma are shown. Curves were compared with the Log-rank test (* *p*-value of α<0.05 was considered significant). (F-G) Mean ± SEM of the plasma viral loads (F) and CD4^+^ T cell counts (G) are shown.

The peak concentrations of VRC01 on the day of the first challenge in the VRC01-only group and in the VRC01-Rh-α_4_β_7_ group were 98.31 ± 25.83µg/ml and 97.66 ± 55.29 µg/ml (mean±SD), respectively (Fig 1B). Of note, for reasons that may be attributed to problems during the infusion, 2 macaques in the VRC01-Rh-α_4_β_7_ group had peak concentrations of VRC01 at the time of the first challenge about 10 fold lower than the mean and below the protective concentration against SHIV_SF162P3_ high-dose challenge ^27,38^. (KT57: 19.145 µg/ml and GH63: 3.350 µg/ml; respectively green and orange in Fig 1B). Thus, these 2 animals were excluded from the SHIV acquisition analysis, but not from other post-infection analysis since VRC01 levels were undetectable in all animals by week 6 post-infection (Fig 1B). Consistent with previous studies, antibodies against VRC01 (ADA) developed around week 3 after infusion in all VRC01-treated animals and levels were similar in both treatment groups (Fig S1). The peak concentration of Rh-α_4_β_7_ in the VRC01-Rh-α_4_β_7_ group at the time of the first challenge was 112±37 µg/ml (mean±SD; Fig 1C). One animal with peak concentrations 10 times higher than the average of the other animals was excluded from the calculation of the average. None of the animals had plasma Rh-α_4_β_7_ concentrations below the expected range and all 9 animals were included in the analysis of the impact of Rh-α_4_β_7_ on acute and early infection parameters. As shown in Fig 1D, all 9 animals in the control group became viremic after 1 to 5 challenges (median of 2 weekly exposures required for infection).

Suboptimal dosing of VRC01 resulted, as expected, in a non-significant delay in SHIV acquisition (Log-rank p=0.074). Nonetheless, the cumulative number of challenges to infect in the VRC01 group was significantly higher than in the control group (Fisher exact test: p=0.02). In the VRC01-Rh-α_4_β_7_ combination treatment, it was possible to detect a significant delay in SHIV acquisition (Log-rank p=0.016; Fig 1E) and the difference with the control group in cumulative number of challenges to infect was highly significant (Fisher exact-test: p<0.001). The median number of challenges needed to infect the two treatment groups was similar (n=4) and the VRC01-Rh-α_4_β_7_ combination did not significantly delay acquisition compared to VRC01 alone (Log-rank p=0.6). Peak plasma viral load (VL) did not significantly differ between the treatment groups and when comparing the treatment groups with the control group (Fig 1F and Fig S2). Set-point VL was ∼1 Log_10_ lower in the VRC01-Rh-α_4_β_7_ group compared with the other 2 groups (Fig 1F). However, the difference did not reach statistical significance (p=0.072 at week 18 p.i.). Confirming previous reports on the effects of Rh-α_4_β_7_ ^23,40^, peripheral CD4^+^ T cells were protected and CD4 counts were significantly higher in the VRC01-Rh-α_4_β_7_ when compared with the other 2 groups (treatment adjusted for time; 2-way Anova p=0.005; Fig 1G). This is at least partially due to Rh-α_4_β_7_-driven lymphocytosis, as we demonstrated in naïve macaques ^22^, and explains the higher baseline CD4 count of the VRC01-Rh-α_4_β_7_ group.

Although gut SIV DNA loads were slightly higher in both treatment groups compared with controls at week 3-4 p.i. (2-3 weeks after 1^st^ virus detection in plasma), by week 20 p.i., VRC01-Rh-α_4_β_7_ treated animals had more than 10-fold lower amounts of cell-associated SIV DNA in the gastrointestinal (GI) tract compared to the controls (Fig 2A). SIV RNA loads in the GI tract of the VRC01-Rh-α_4_β_7_ treated animals were also 100 times lower, on average, than in the control group during the post-acute phase (week 7-8 p.i.; Fig 2B). However, by week 20 p.i. several animals in the control group had undetectable SIV-RNA levels in the GI tract and the difference with the VRC01-Rh-α_4_β_7_ lost its significance. Interestingly, no significant differences in SIV loads were found by directly comparing the VRC01-Rh-α_4_β_7_ combination group and the VRC01-alone group and the SIV loads in the VRC01-only group often averaged between the other 2 groups. Vaginal SIV DNA and RNA loads did not differ in the acute nor in the early chronic phase of the infection among the treatment groups (Fig S3). Finally, although the SIV-DNA loads at necropsy (around 20 weeks p.i.) were on average slightly lower in the jejunum and iliac lymph nodes of the VRC01-Rh-α_4_β_7_ group compared with the other 2 groups, the differences were not significant (Fig S4). No differences were detected in other tissues and lymph nodes at necropsy (Fig S4).

**Fig 2.**
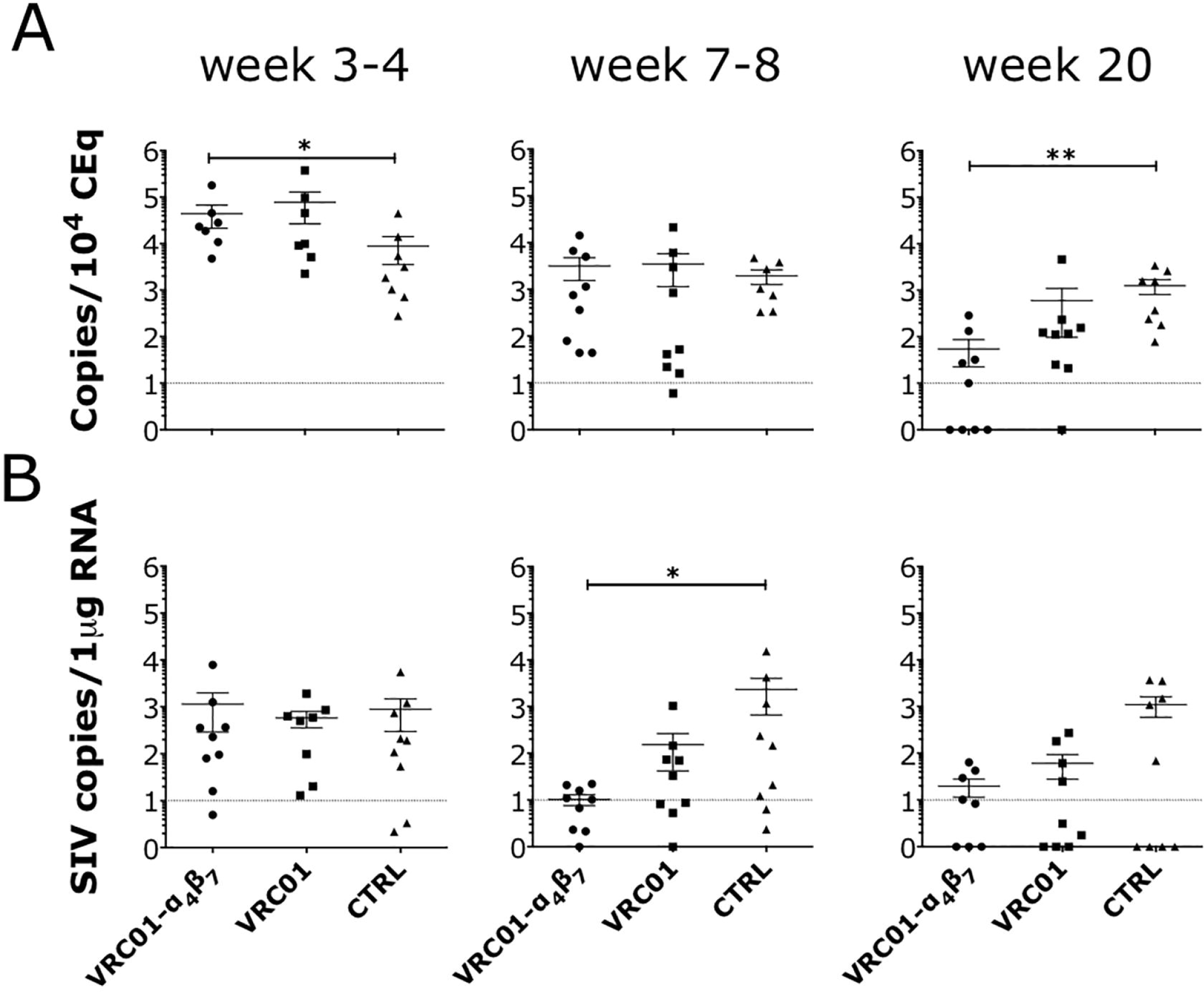
The Rh-α_4_β_7_-VRC01 combination reduces viral DNA and RNA in the gut. Copies of SIV DNA (A) and RNA (B) from colorectal biopsies at the indicated times after infection were quantified by gag-qPCR (normalized on albumin content) and by RT-qPCR (normalized on RNA content) respectively. The dotted line indicates the lower limit of detection (LLOD) of the assay. Data from the treatment groups were compared with the control by Kruskal-Wallis test and the results of the Dunn’s multiple comparisons post-hoc test are shown (*p*-value of * α<0.05 and ** α <0.01 were considered significant).

### VRC01-Rh-α_4_β_7_ decreases IL-17-producing cells in acute infection and increases IFNγ-producing cells in the chronic phase

Mononuclear cells isolated from blood and colorectal biopsies in the acute phase of infection (week 3-4 p.i.) and from ileum, jejunum and colorectal tissue at necropsy (chronic phase) were stimulated with PMA/ionomycin to determine the frequency of IL-17 (acute and chronic phase samples) and IFN*γ*, IL-2, TNF*α* and IL21-producing cell subsets (chronic phase samples). VRC01-Rh-α_4_β_7_ treated animals had significantly lower frequencies of IL-17 producing CD8^+^ T cells in blood and of IL-17 producing CD4^+^ T cells and CD3^-^ NKp44^+^ NK cells in the colorectal tissue compared to the control group (Fig 3A-B). For these subsets, the difference did not reach significance when compared with the VRC01-only group. However, the frequency of IL-17-producing NKG2A^+^ NK cells in blood was significantly lower in the VRC01-Rh-α_4_β_7_ group compared with the VRC01-only group (Fig 3A). In contrast, in the chronic phase, the frequency of IFN*γ*-producing CD4^-^ T cells was higher in the VRC01-Rh-α_4_β_7_ group compared with the control group in the small intestine (Fig 3C). No significant differences in IFN*γ*-producing cells were noted in blood or upon direct comparison of the VRC01-only group with the VRC01-Rh-α_4_β_7_ group nor between the VRC01-only group and the control. No significant differences in IL-17, IL-2, TNF*α* and IL21-producing cells were noted in these tissues in the chronic phase of infection.

**Fig 3.**
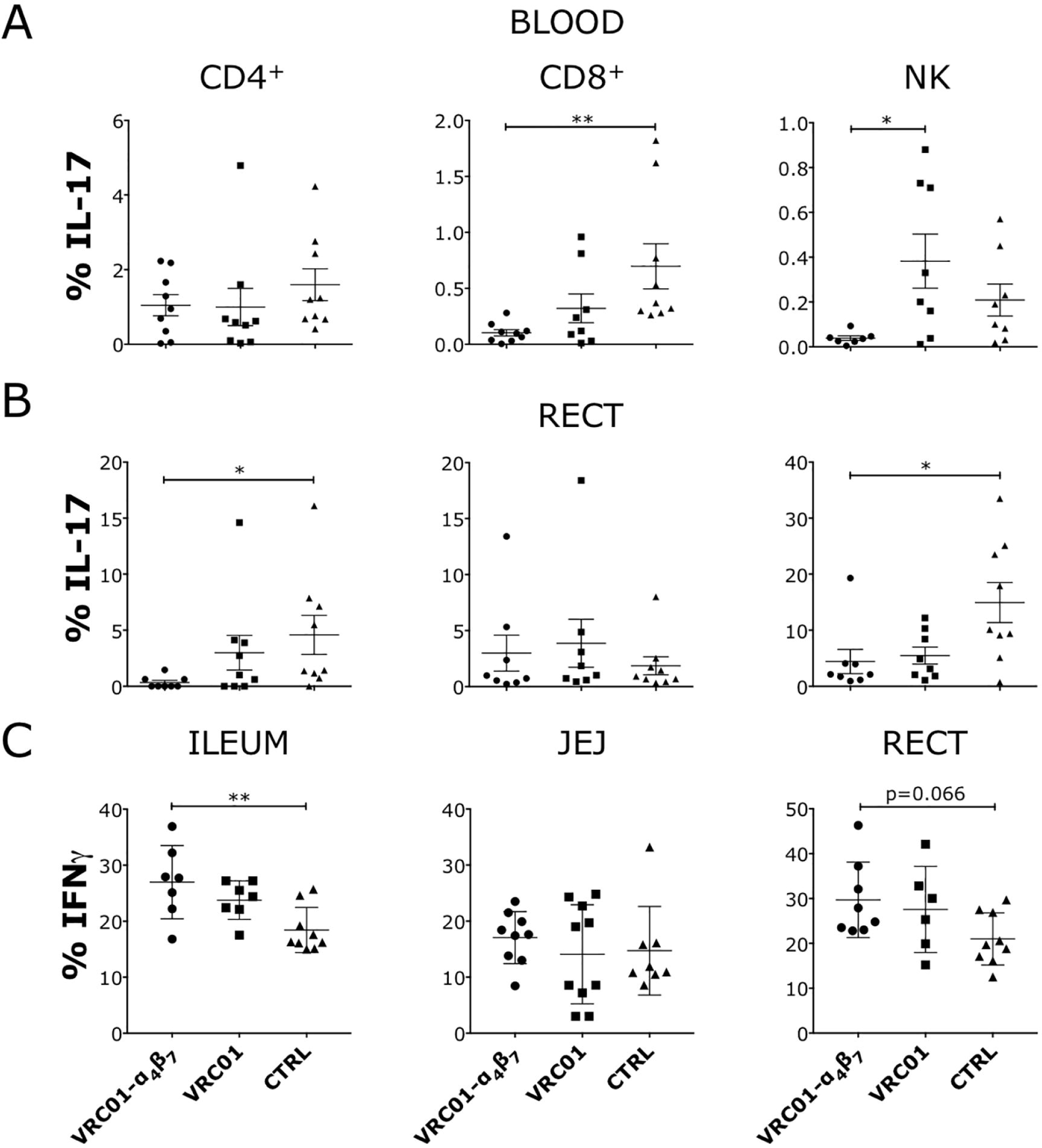
Decreased IL-17 and increased IFN□ producing cells in the gut of Rh-α_4_β_7_-VRC01-treated macaques. (A-B) The frequency of IL-17-secreting cells within the indicated subsets in blood (A) and colorectal biopsies (B) collected 2 weeks after the first detection of virus in plasma (3-4 weeks post-infection) are shown. (C) The frequency of IFN-□-secreting cells within CD8^+^ T cells in the indicated tissues are shown. (A-C) NK-like cells were defined as CD3^-^NKG2A^+^ in the blood and CD3^-^NKp44^+^ in the colorectal tissue. Data from the treatment groups were compared with the control by Kruskal-Wallis test and the results of the Dunn’s multiple comparisons post-hoc test and the Mann-Whitney test to compare the treatment groups between each other are shown (*p*-value of * α<0.05 and ** α<0.01 were considered significant).

Interestingly, when we investigated how the treatments had impacted immune cells in the lymph nodes at necropsy, we found that in the VRC01-only group had a lower frequency of CD25^+^ T cells (both CD4^+^ and CD4^-^, Fig 4) compared with the control group. Rh-*α*_4_*β*_7_ in the VRC01-Rh-*α*_4_*β*_7_ may have interfered with this decrease since the difference was not significant between the VRC01-Rh-*α*_4_*β*_7_ group and the controls. Moreover, we found that the VRC01-treated animals had lower frequencies of CXCR3 CCR6 double positive CD4^-^ and CD4^+^ T cells compared with both the VRC01-Rh-*α*_4_*β*_7_ and the control groups (Fig 4A and B). Finally, the VRC01-only treated animals had lower CXCR5^+^ CD4^+^ follicular T cells in the lymph nodes compared with the controls (Fig 4A). No other differences were noted among CD103^+^, CD69^+^, and Treg-like CD127^-^CD25^+^ cells within the CD4^+^ or CD4^-^ cell subsets in the lymph nodes between the groups (not shown). Phenotyping of mononuclear cells isolated from blood during the chronic phase of infection showed that Rh-α_4_β_7_ treatment was associated with an increase of CCR6^+^CD95^-^ CD4^-^ cells compared to both control groups (Fig S5).

**Fig 4.**
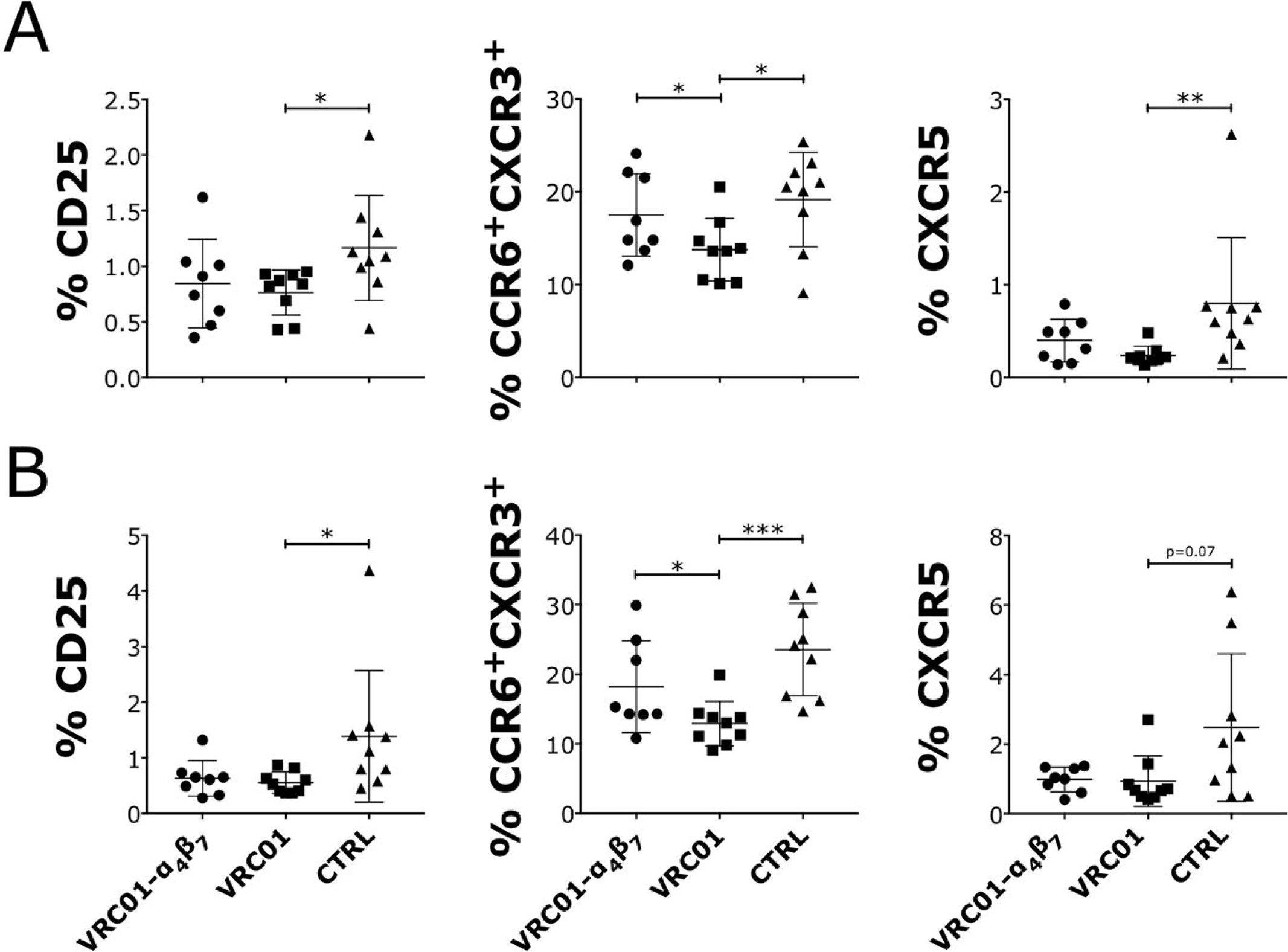
Decreased frequencies of CD25^+^, CCR6^+^ CXCR3^+^ and CXCR5^+^ T cells in the lymph nodes of VRC01 pretreated, SHIV-infected macaques. (A-B) At necropsy, cells were isolated from inguinal lymph nodes and analyzed by flow cytometry. The frequencies of subsets within the CD4^+^ (A) and CD8^+^ (B) T cells that were significantly different in any of the treatment groups compared to the controls by Kruskal-Wallis test are shown. The results of the Dunn’s multiple comparisons post-hoc test and the Mann-Whitney test to compare the treatment groups between each other are shown (*p*-value of * α<0.05, ** α<0.01 and *** α<0.001 were considered significant).

### T cell responses in the VRC01-Rh-α_4_β_7_ are primarily directed against the V2-loop of the SHIV_AD8-EO_ envelope protein

Blood T cell responses were analyzed against pooled 15-mer overlapping peptides from the consensus B envelop and gag proteins around week 18 post-infection. Interestingly, T cell responses against the envelope peptides in the VRC01-Rh-α_4_β_7_ group were virtually undetectable and significantly lower than the T cell responses in the control group (Fig S6). T cell responses in the VRC01 group were more similar to the control group, but the difference between the VRC01-only and VRC01-Rh-α_4_β_7_ groups reached statistical significance only for TNF*α* and IL22-secreating CD4^+^ T cells (Fig S6A). The responses to gag peptides were generally lower than those to the envelope peptides at this stage of infection in all groups and differences between groups could not be determined.

Since the sequence of the consensus B envelope differs substantially from the sequence of the SHIV_AD8-EO_ envelope in the V1V2 region (Fig S7) and a specific response against the V2-loop was found in the Rh-α_4_β_7_-treated animals of the cART-Rh-α_4_β_7_ study ^17^, we synthetized seven 20-mer, 14aa overlapping peptides spanning the V1V2 region that differs between the SHIV_AD8-EO_ and the consensus B envelope and used them to probe T cells and antibody responses. Interestingly, the VRC01-Rh-α_4_β_7_ macaques had higher frequencies of IFN*γ*-producing CD4^+^ T cells in response to the V1V2 peptide pool than both the VRC01 and control groups (Fig 5A). In contrast, TNF*α*-producing CD4^+^ T cells and IFN-*γ*-producing CD8^+^ T cells were higher in both treatment groups compared with the control group (Fig 5A-B). A higher IL-17 response in the CD8^+^ T cell subset was also noted in the VRC01-Rh-α_4_β_7_ group compared to the controls (Fig 5B). Finally, a higher frequency of TNF-*α*-CD8^+^ T cells was present in the VRC01-Rh-α_4_β_7_ group compared with the VRC01-only group. In summary, higher T cell responses were detected against the V1V2 loop in the VRC01-Rh-α_4_β_7_ group particularly compared to control animals. Interestingly, the frequency of CD8^+^ T cells secreting TNF-*α* in response to the V1V2 peptides inversely correlated with viral load (Fig S8), suggesting that a significant role is played by V1V2 responses in controlling viral replication.

**Fig 5.**
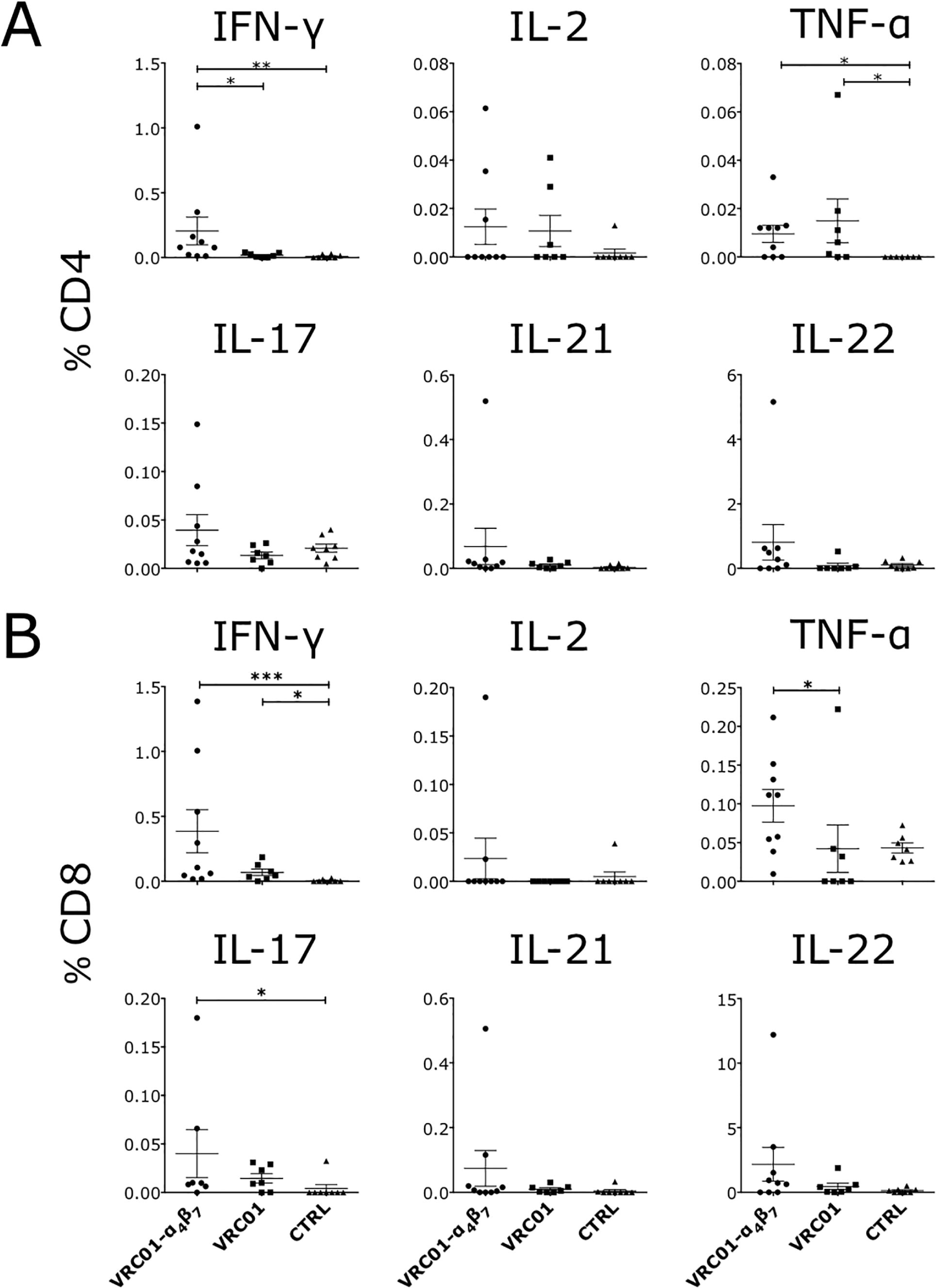
T cell responses to the SHIV_AD8-EO_ V2-loop peptides are higher in Rh-α_4_β_7_-VRC01 treated macaques. PBMCs isolated around 18 weeks post infection were stimulated with a pool of 7 20mers with 14aa overlap peptides spanning the V2 loop of the SHIV_AD8-EO_ envelope for 5hrs. The frequency of cells secreting the indicated cytokines are shown for the CD4^+^ and CD8^+^ T cell subsets after subtraction of the baseline values (in absence of peptides). The results of the Dunn’s multiple comparisons post-hoc test (after the Kruskal-Wallis test controlled for multiple comparisons) and the Mann-Whitney test to compare the treatment groups between each other are shown (*p*-value of * α<0.05, ** α<0.01 and *** α<0.001 were considered significant).

Total anti-HIV envelope antibodies were tested against the HIV-Bal envelope protein and no differences were noted between the groups. Moreover, a peptide scan against consensus B envelope peptides (with the 8 peptides corresponding to the V1V2 loop replaced by the 7 SHIV-AD8-specific peptides that we had synthetized) was carried out on sera from 4 animals in each group with the highest antibody responses. No clear differences in the response to specific regions of the envelope were noted between the groups (Fig S8).

## DISCUSSION

bNAbs are being tested in the clinic for the prevention and therapy of HIV infection. VRC01 is the first to reach efficacy testing and other bNAbs will soon follow ^41,42^. However, it is clear that individual bNAbs cannot be used alone as a single intervention. Combinations of more bNAbs or bi- or tri-specific molecules need to be employed to achieve better and more durable protection from HIV acquisition ^33,43-45^. Moreover, recent data suggest that bNAbs treatment may impact immune responses to infection ^46-48^. This feature represents a potential new therapeutic approach toward an HIV-1 cure. Rh-α_4_β_7_ has also demonstrated the ability to partially prevent SIV infection in macaques ^16^ and treatment of SIV infected macaques with Rh-α_4_β_7_ in combination with cART has shown its potential utility in inducing long-term control of SIV replication without eradicating the virus ^17^. Nonetheless, much more needs to be understood about the ability of bNAbs and α_4_β_7_-blockage to impact immune responses against SIV/HIV.

The present study represents the first investigation of the combination of a bNAb and Rh-α_4_β_7_. We aimed to determine how this dual-treatment might alter key features of acute and early-chronic infection, including anti-viral immune responses. In order to observe such effects beyond the powerful protective activity of VRC01, we performed the study in a setting of suboptimal amounts of VRC01 against repeated challenges with a very high viral inoculum of a pathogenic SHIV.

Overall, we determined that Rh-α_4_β_7_ has the potential to increase the protective activity of suboptimal dosing of VRC01. However, larger studies with a more physiologically relevant, low-dose viral challenge will be needed to precisely quantify the incremental effect of combining Rh-α_4_β_7_ and VRC01 over VRC01 on SHIV acquisition. Interestingly, we were able to determine that some of the key effects of the Rh-α_4_β_7_ that were previously described ^16,23,40^ are retained in the SHIV model. They include the ability of the Rh-α_4_β_7_ to protect circulating CD4^+^ T cells, a modest effect on the viral set-point and a profound effect on the gut viral load. However, we did not find a decrease viral load in any other tissue or lymph nodes as it was described in the SIV model ^23^.

Interestingly, most of the differences that we noted are significant only when the VRC01-Rh-α_4_β_7_ is compared with the control group and the data from the VRC01-only group fell in between the other 2 groups. This suggests that the presence of minimal quantities of VRC01 at the time of infection may have facilitated the impact of Rh-α_4_β_7_ on virologic and immune parameters, hinting at the possibility of a synergistic effect of the two antibodies. This, also, will need further investigation.

Notably, we found that during the acute phase of infection (which coincided with the Rh-α_4_β_7_-treatment) the VRC01-Rh-α_4_β_7_ group had a lower frequency of IL-17 producing T cells in the gut. This may be due to the profound impact of Rh-α_4_β_7_ on CCR6^+^ T cells that we noted in absence of infection ^22^ and may help explain the overall decrease in T cell responses that was observed in the VRC01-Rh-α_4_β_7_ group when pools for the entire envelope and gag proteins were used. Perhaps because the Rh-α_4_β_7_ was administered during the earliest stages of the infection, we did not see an impact on the antibody responses as was described in the ART-Rh-α_4_β_7_ combination study ^17^. However, treatment with Rh-α_4_β_7_ increased the T cell responses against the V2 region of the envelope and our data suggest a role for these responses in maintaining virologic control. How and why this happens requires further investigation. It is possible to speculate that in the presence of Rh-α_4_β_7_ immune cell priming occurs more efficiently outside the mucosal tissues and/or that the Rh-α_4_β_7_ changes the antigenicity of the envelope or antigen processing.

Interestingly, the only VRC01-specific effect we observed appeared in the lymph nodes, where in the VRC01-only treated animals we detected significantly lower frequencies of CD25^+^ and CXCR5^+^ CD4^+^ T cells. In the VRC01-Rh-α_4_β_7_ group this effect was still present, but less pronounced. This further supports the published observations that suggest a direct impact of bNAbs on the antiviral immune response ^47,49^. Nonetheless, the amount of time the animals were exposed to meaningful concentrations of VRC01 during infection was very short and the data should be interpreted with caution. Importantly, the VRC01-Rh-α_4_β_7_ group had higher frequencies of IFN*γ* producing T cells in both the large and small intestine. This increase suggests a protective activity of the Rh-α_4_β_7_ on the gut immune system and helps explain the long-term survival of SIV infected animals treated with Rh-α_4_β_7_ during acute infection (Aftab Ansari, unpublished and reported at CROI 2018 by J. Arthos). Boosting effective mucosal immune responses may also contribute to explain the virologic control seen when the Rh-α_4_β_7_ was used in combination with cART as therapeutic approach in Byrareddy et. al, Science 2016 ^17^. This control was not replicated in other similar studies (as reported by Di Mascio and Fauci at AIDS2018). Nonetheless, our study adds to the considerable amount of work that supports a profound effect of the Rh-α_4_β_7_ on immune cells and the immune responses.

In conclusion, a suboptimal dose of VRC01 in combination with the Rh-α_4_β_7_ significantly delayed infection and impacted the availability and distribution of immune cell subsets as well as the T cell responses to the virus. VRC01 and Rh-α_4_β_7_ impact HIV infection by distinct mechanisms, neither of which is fully understood. This study gives the first insight into the combination of these antibodies and contributes to our understanding their effect on HIV infection. Future studies should address how other bNAbs combinations with Rh-α_4_β_7_ could be harnessed in different therapeutic and preventive settings to fight HIV infection.

## MATERIALS AND METHODS

### Macaques and treatments

A total of 27 adult female Indian rhesus macaques (average weight 8.1Kg, range: 4.5, 13.05 and age 9.5 years, range: 3.9, 18) were socially housed, indoors, in climate controlled conditions at the Tulane National Primate Research Center (TNPRC) in compliance with the regulations under the Animal Welfare Act, the Guide for the Care and Use of Laboratory Animals ^50^. All procedures were approved by the Animal Care and Use Committee of the TNPRC and in compliance with animal care procedures. The animals were euthanized at the end of the study using methods consistent with recommendations of the American Veterinary Medical Association (AVMA) Panel on euthanasia and per the recommendations of the IACUC.

Macaques were divided in three groups of 9 animals each. Animals were administered intravenously 1) 1 injection of 10mg/Kg of VRC01 + 1 injection of 25mg/Kg of Rh-α_4_β_7_ mAb; followed by 5 additional injections of 25mg/Kg of Rh-α_4_β_7_ mAb every 3 weeks 2) 1 injection of 10mg/Kg of VRC01; 3) 1 injection of 10mg/Kg of control Human IgG and 25mg/Kg of control Rhesus IgG. Starting 3 days after treatment, macaques were challenged weekly with intravaginal administration of SHIV_AD8-OE_ (1000 TCID_50_/challenge) for 8 weeks. Blood was collected weekly to monitor infection as described below. After two consecutive positive SIV PCRs (3-4 weeks post infection; acute phase) fresh rectal biopsies and blood were used for cell isolation as described below. Subsequently, blood was collected every two weeks throughout the study. Rectal and vaginal samples were also collected at 7-8 weeks post infection (post-acute phase). At necropsy, blood, lymph nodes, gut, brain, vaginal and cervical tissues were harvested and used for cell isolation.

### Measurement of plasma levels of VRC01 and Rh-α_4_β_7_

Levels of VRC01 were measured as described in ^27^. Levels of rhesus Rh-α_4_β_7_ antibody in macaque plasma were measured using the α_4_β_7_-expressing human T cell line HuT-78 (NIH AIDS Reagent Program, Division of AIDS, NIAID, NIH: HUT 78 from Dr. Robert Gallo) in a flow cytometry-based assay as described in (Calenda 2018, Byrareddy 2014) using the standard curve method. Briefly, HuT78 cells were first incubated for 2-3 days in complete RPMI 1640 media containing 100nM retinoic acid to increase the surface expression of α_4_β_7._ Cells (150,000/condition) were stained with LIVE/DEAD Aqua dye (Thermo Fisher Scientific, Waltham, MA) for live/dead discrimination, incubated for 30min at 4°C with the plasma to be tested (1:10 diluted in PBS) obtained from macaques from the VRC01-α_4_β_7_ treatment group before (baseline, BL) and up to 6 weeks after treatment. Cells were then washed and incubated for 30 min at 4°C with anti-rhesus IgG1 (NHP resource center, antibody 7H11, in house biotinylated with EZ-link NHS-biotin (Thermo Fisher Scientific) following the manufacturer’s instructions), washed again and resuspended in neutravidin-PE (Thermo Fisher Scientific) for 20 min at 4°C. PE fluorescence was analyzed on a flow cytometer. For the standard curve, baseline plasma was pooled and spiked with serial dilutions of Rh-α_4_β_7_ (2500 µg/ml – 0 µg/ml).

### Anti-VRC01 and anti-Rh-α_4_β_7_ antibodies

Levels of anti-VRC01 antibodies were measured as described in ^32^. Levels of anti-Rh-α_4_β_7_ antibodies were measured via lamda light chain detection assay. ELISA plates were coated with Rh-α_4_β_7_ (10µg/ml) overnight at 4°C, washed and blocked with TBS 2% BSA 0.1% Tween20 for 2 hours at room temperature (RT). Test plasma was serially diluted in dilution buffer (starting at 1:10 then serial 1:4 dilutions), 100 µl applied to the plates and incubated 1 hour at RT. Plates were washed and incubated with anti-Ig human lambda light chain-biotin (Miltenyi), which does not recognize the kappa chain of the Rh-α_4_β_7_ 1 hour at RT. Plates were washed and incubated with diluted streptavidin-HRP (Invitrogen) 1 hour at RT. Plates were washed and enzymatic activity detected by adding TMB substrate and read on a luminometer at 450nm. Endpoint was the highest dilution with OD 2-fold higher the pre-treatment sample.

### SHIV_AD8-EO_ stock generation and titration

The full-length proviral plasmid pSHIVAD8-EO was a gift from Dr. Malcom Martin. Virus stocks were prepared by transfecting 293T cells with 5µg of the pSHIV_AD8-EO_ molecular clone using Lipofectamine 2000 (Invitrogen). Culture supernatant was collected 60 hours later, clarified by centrifugation (300g 10 minutes, 4°C) and used to infect CD8-depleted, PHA-activated rhesus macaque PBMC. Cells were incubated overnight in 293T supernatants, washed and resuspended in RPMI 10% FBS medium for 10 days. Supernatants from parallel cultures were pooled on day 7, clarified by centrifugation (10,000g, 15minutes, 4°C), aliquoted, and stored at −80°C. The resulting stock was titrated in PHA-activated rhesus macaque PBMC.

### SIV viral loads

Macaque infection was confirmed by SIVgag nested PCR on PBMC as described [49]. Plasma samples were obtained from EDTA-treated whole blood and used for the determination of plasma VL by SIVgag qRT-PCR [50] (quantitative Molecular Diagnostics Core, AIDS and Cancer Virus Program Frederick National Laboratory). DNA and RNA were extracted from snap frozen tissues using DNeasy/RNeasy blood and tissue kits (Qiagen) following the manufacturer’s instructions. Tissue viral DNA loads were quantified using the standard curve method and normalized by albumin copy numbers by gag-qPCR as described in ^51^. For tissue RNA loads, 1µg of total RNA was retrotranscribed to DNA using the VILO Kit (Thermo Fisher) quantified by gag-qPCR ^51^.

### Cell isolation and flow cytometry

3-4 weeks post infection, PBMCs were isolated using Ficoll-Hypaque density gradient centrifugation and cells from rectal biopsies were isolated by enzymatic digestion in HBSS containing 2mg/mL Collagenase IV (Worthington Biochemical) and 1mg/ml of Human Serum Albumin (Sigma-Aldrich), shaking at 37°C for 50 minutes. The resulting cell suspension was passed through a 40μm cell strainer and washed with PBS. PBMCs and rectal cells were then stimulated with 60ng/ml PMA, 0.5μg/ml Ionomycin and 5μg/ml Brefeldin A (BFA) for 4 hours at 37°C, stained with LIVE/DEAD Aqua viability dye (Thermo Fisher Scientific) and incubated with a cocktail of different panels of monoclonal antibodies as listed in the tables in Fig S9.

At the time of necropsy, PBMCs were isolated by Ficoll-Hypaque density gradient centrifugation. Enzymatic digestion was used to isolate cells from jejunum, ileum, colorectal, vaginal and cervical tissues as described above. Spleen and LNs (axillary, mesenteric, inguinal, iliac) were cut in small pieces and passed directly through a 40μm cell strainer. Isolated cells were washed, frozen for phenotyping and stimulation experiments at a later time. For PBMC stimulation experiments, up to 3×10^6^ cells/sample were thawed, plated on a plate pre-coated with 2.5μg/ml goat anti mouse (GAM) IgGs and cross linked with 10μg/ml anti-CD28 and anti-CD49d antibodies (Sigma Aldrich). Cells were stimulated either with SIVMAC239 GAG peptide pool (1μg/ml; 125 15mers with 11 aa overlap, AIDS reagents program, Division of AIDS, NIAID, NIH), HIV-1 Consensus B Env Peptide Set (2μg/ml; 211 15mers with 11 aa overlap, AIDS reagents program, Division of AIDS, NIAID, NIH) or a SHIV_AD8OE_ specific Env V1-V2 peptide pool (2μg/condition; 7 20mers with 14 aa overlap, Peptide 2.0, Chantilly, VA). Pooled cells stimulated with 1μg/ml PMA/Ionomycin were used a positive activation control. One hour later 10μg/ml BFA and monensin (GolgiStop, BD Biosciences) were added to each well. After 5 hours, cells were transferred to a FACS plate and stained with the panels listed in table 1. To maximize CD107a detection, antibody staining was performed during stimulation. For gut tissue stimulation experiments, cells isolated from colorectum, jejenum and ileum where thawed and stimulated with 60ng/ml PMA, 0.5μg/ml Ionomycin and 5μg/ml Brefeldin A (BFA) for 4 hours at 37°C, stained with LIVE/DEAD Aqua viability dye (Thermo Fisher Scientific) and incubated with a cocktail of different panels of monoclonal antibodies as listed in the tables in Fig S9. For phenotyping experiments, cells were thawed and stained with the panels listed in tables in Fig S9.

### Pepscan

Peptide scan was performed against consensus B envelope peptides with the 8 peptides corresponding to the V1-V2 loop replaced by the 7 SHIV_AD8OE_-specific peptides that we had synthetized was performed on serum samples from the 4 animals from each group showing the highest antibody response by ABL Inc.

### Statistics

To analyze the differences in SHIV acquisition, the survival curves generated with time to first viral detection in plasma from each treatment group were compared to each other directly and each with the curve from the control group using the Log-rank (Mantel-Cox) test. The cumulative number of challenges needed to infect in each group was also compared to the cumulative number in each other group by Fisher exact test. Viral loads and CD4 counts were compared by two-way ANOVA for repeated measures. To address whether either treatment had a significantly different effect on infection and immunological parameters than the control group, data was analyzed using Kruskal-Wallis non-parametric test adjusted for multiple comparisons followed by the Dunn’s multiple comparisons post-hoc test. To address whether treatment groups differed from each other a Mann-Whitney unpaired t-test was performed. All analyses were performed using the GraphPad Prism software V7. Significant *p*-values of α<0.05 (*), α<0.01 (**) and α <0.001 (***) are indicated.

## Supporting information

## Acknowledgments

This work was supported by NIAID grant R01AI098546-05. We kindly thank Dr. Aftab A. Ansari for the many scientific discussions on the effect of the Rh-α_4_β_7_ and steadfast support for our work. We thank Dr. Keith Reimann and Mr. Adam Busby (Mass Biologics; supported by grants AI126683 and OD010976) for the preparation of the Rh-α_4_β_7_ and the Rh-IgG and for the assistance during the studies. We thank Dr. Malcom Martin and his team (Laboratory of Molecular Microbiology at NIAID, NIH) for providing the SHIV_AD8-EO_ plasmid and related protocols. We are very grateful to Jeffrey Lifson, Rebecca Shoemaker and the rest of the team from the AIDS and Cancer Virus Program (Leidos Biomedical Research, Inc.) for supporting this work and being always timely in measuring the plasma viral load. Finally, we acknowledge the outstanding help and assistance of the professional staff, animal technicians and caretakers of the Tulane National Primate Research Center and the staff of the Population Council and Rockefeller University resource centers for their assistance and help in the conduction of the studies presented in this report.

## Author contributions

EM, JA, conceived and designed the study; GC, IF, GAB coordinated the receipt and processing of macaque samples, run the assays and analyzed the data; AP and KW measured VRC01 concentrations and anti-VRC01 antibodies; SM processed samples and run virologic assays; BJ, JLB and AG coordinated the macaque studies, sample collection and animal welfare; EM and KR performed statistical analysis; EM, JA, CC, ASF, JRM interpreted the data and wrote the manuscript.

## References

1. Schweighoffer T, Tanaka Y, Tidswell M, et al. Selective expression of integrin alpha 4 beta 7 on a subset of human CD4+ memory T cells with Hallmarks of gut-trophism. J Immunol. 1993;151(2):717–729.

2. Reese SR, Kudsk KA, Genton L, Ikeda S. l-selectin and alpha4beta7 integrin, but not ICAM-1, regulate lymphocyte distribution in gut-associated lymphoid tissue of mice. Surgery. 2005;137(2):209–215.

3. Meenan J, Spaans J, Grool TA, Pals ST, Tytgat GN, van Deventer SJ. Altered expression of alpha 4 beta 7, a gut homing integrin, by circulating and mucosal T cells in colonic mucosal inflammation. Gut. 1997;40(2):241–246.

4. Brenchley JM, Schacker TW, Ruff LE, et al. CD4+ T cell depletion during all stages of HIV disease occurs predominantly in the gastrointestinal tract. J Exp Med. 2004;200(6):749– 759.

5. Guadalupe M, Reay E, Sankaran S, et al. Severe CD4+ T-cell depletion in gut lymphoid tissue during primary human immunodeficiency virus type 1 infection and substantial delay in restoration following highly active antiretroviral therapy. J Virol. 2003;77(21):11708– 11717.

6. Mehandru S, Poles MA, Tenner-Racz K, et al. Primary HIV-1 infection is associated with preferential depletion of CD4+ T lymphocytes from effector sites in the gastrointestinal tract. J Exp Med. 2004;200(6):761–770.

7. Li Q, Duan L, Estes JD, et al. Peak SIV replication in resting memory CD4+ T cells depletes gut lamina propria CD4+ T cells. Nature. 2005;434(7037):1148–1152.

8. Cicala C, Martinelli E, McNally JP, et al. The integrin alpha4beta7 forms a complex with cell-surface CD4 and defines a T-cell subset that is highly susceptible to infection by HIV-1. Proc Natl Acad Sci U S A. 2009;106(49):20877–20882.

9. Kader M, Wang X, Piatak M, et al. Alpha4(+)beta7(hi)CD4(+) memory T cells harbor most Th-17 cells and are preferentially infected during acute SIV infection. Mucosal Immunol. 2009;2(5):439–449.

10. Wang X, Xu H, Gill AF, et al. Monitoring alpha4beta7 integrin expression on circulating CD4+ T cells as a surrogate marker for tracking intestinal CD4+ T-cell loss in SIV infection. Mucosal Immunol. 2009;2(6):518–526.

11. Sivro A, Schuetz A, Sheward D, et al. Integrin alpha4beta7 expression on peripheral blood CD4(+) T cells predicts HIV acquisition and disease progression outcomes. Sci Transl Med. 2018;10(425).

12. Martinelli E, Veglia F, Goode D, et al. The frequency of alpha4beta7high memory CD4+ T cells correlates with susceptibility to rectal SIV infection. J Acquir Immune Defic Syndr. 2013.

13. Martinelli E, Tharinger H, Frank I, et al. HSV-2 infection of dendritic cells amplifies a highly susceptible HIV-1 cell target. PLoS Pathog. 2011;7(6):e1002109.

14. Goode D, Truong R, Villegas G, et al. HSV-2-driven increase in the expression of alpha4beta7 correlates with increased susceptibility to vaginal SHIV(SF162P3) infection. PLoS Pathog. 2014;10(12):e1004567.

15. Shannon B, Yi TJ, Thomas-Pavanel J, et al. Impact of Asymptomatic Herpes Simplex Virus Type 2 Infection on Mucosal Homing and Immune Cell Subsets in the Blood and Female Genital Tract. The Journal of Immunology. 2014;192(11):5074–5082.

16. Byrareddy SN, Kallam B, Arthos J, et al. Targeting alpha4beta7 integrin reduces mucosal transmission of simian immunodeficiency virus and protects gut-associated lymphoid tissue from infection. Nat Med. 2014;20(12):1397–1400.

17. Byrareddy SN, Arthos J, Cicala C, et al. Sustained virologic control in SIV+ macaques after antiretroviral and alpha4beta7 antibody therapy. Science. 2016;354(6309):197–202.

18. Joag VR, McKinnon LR, Liu J, et al. Identification of preferential CD4+ T-cell targets for HIV infection in the cervix. Mucosal Immunol. 2016;9(1):1–12.

19. Parrish NF, Wilen CB, Banks LB, et al. Transmitted/founder and chronic subtype C HIV-1 use CD4 and CCR5 receptors with equal efficiency and are not inhibited by blocking the integrin alpha4beta7. PLoS Pathog. 2012;8(5):e1002686.

20. Nawaz F, Goes LR, Ray JC, et al. MAdCAM costimulation through Integrin-alpha4beta7 promotes HIV replication. Mucosal Immunol. 2018.

21. Arrode-Bruses G, Goode D, Kleinbeck K, et al. A Small Molecule, Which Competes with MAdCAM-1, Activates Integrin alpha4beta7 and Fails to Prevent Mucosal Transmission of SHIV-SF162P3. PLoS Pathog. 2016;12(6):e1005720.

22. Calenda G, Keawvichit R, Arrode-Bruses G, et al. Integrin alpha4beta7 Blockade Preferentially Impacts CCR6(+) Lymphocyte Subsets in Blood and Mucosal Tissues of Naive Rhesus Macaques. J Immunol. 2018;200(2):810–820.

23. Santangelo PJ, Cicala C, Byrareddy SN, et al. Early treatment of SIV+ macaques with an alpha4beta7 mAb alters virus distribution and preserves CD4(+) T cells in later stages of infection. Mucosal Immunol. 2017.

24. Baba TW, Liska V, Hofmann-Lehmann R, et al. Human neutralizing monoclonal antibodies of the IgG1 subtype protect against mucosal simian-human immunodeficiency virus infection. Nat Med. 2000;6(2):200–206.

25. Mascola JR, Stiegler G, VanCott TC, et al. Protection of macaques against vaginal transmission of a pathogenic HIV-1/SIV chimeric virus by passive infusion of neutralizing antibodies. Nat Med. 2000;6(2):207–210.

26. Parren PW, Marx PA, Hessell AJ, et al. Antibody Protects Macaques against Vaginal Challenge with a Pathogenic R5 Simian/Human Immunodeficiency Virus at Serum Levels Giving Complete Neutralization In Vitro. J Virol. 2001;75(17):8340–8347.

27. Pegu A, Yang ZY, Boyington JC, et al. Neutralizing antibodies to HIV-1 envelope protect more effectively in vivo than those to the CD4 receptor. Sci Transl Med. 2014;6(243):243ra288.

28. Anderson DJ, Politch JA, Zeitlin L, et al. Systemic and topical use of monoclonal antibodies to prevent the sexual transmission of HIV. AIDS. 2017;31(11):1505–1517.

29. Zhou T, Georgiev I, Wu X, et al. Structural basis for broad and potent neutralization of HIV-1 by antibody VRC01. Science. 2010;329(5993):811–817.

30. Wu X, Yang ZY, Li Y, et al. Rational design of envelope identifies broadly neutralizing human monoclonal antibodies to HIV-1. Science. 2010;329(5993):856–861.

31. Gautam R, Nishimura Y, Pegu A, et al. A single injection of anti-HIV-1 antibodies protects against repeated SHIV challenges. Nature. 2016;533(7601):105–109.

32. Ko SY, Pegu A, Rudicell RS, et al. Enhanced neonatal Fc receptor function improves protection against primate SHIV infection. Nature. 2014;514(7524):642–645.

33. Xu L, Pegu A, Rao E, et al. Trispecific broadly neutralizing HIV antibodies mediate potent SHIV protection in macaques. Science. 2017;358(6359):85–90.

34. Steinhardt JJ, Guenaga J, Turner HL, et al. Rational design of a trispecific antibody targeting the HIV-1 Env with elevated anti-viral activity. Nat Commun. 2018;9(1):877.

35. Bournazos S, Gazumyan A, Seaman MS, Nussenzweig MC, Ravetch JV. Bispecific Anti-HIV-1 Antibodies with Enhanced Breadth and Potency. Cell. 2016;165(7):1609–1620.

36. Nishimura Y, Shingai M, Willey R, et al. Generation of the pathogenic R5-tropic simian/human immunodeficiency virus SHIVAD8 by serial passaging in rhesus macaques. J Virol. 2010;84(9):4769–4781.

37. Gautam R, Nishimura Y, Lee WR, et al. Pathogenicity and mucosal transmissibility of the R5-tropic simian/human immunodeficiency virus SHIV(AD8) in rhesus macaques: implications for use in vaccine studies. J Virol. 2012;86(16):8516–8526.

38. Rudicell RS, Kwon YD, Ko SY, et al. Enhanced potency of a broadly neutralizing HIV-1 antibody in vitro improves protection against lentiviral infection in vivo. J Virol. 2014;88(21):12669–12682.

39. Kwa S, Kannanganat S, Nigam P, et al. Plasmacytoid dendritic cells are recruited to the colorectum and contribute to immune activation during pathogenic SIV infection in rhesus macaques. Blood. 2011;118(10):2763–2773.

40. Ansari AA, Reimann KA, Mayne AE, et al. Blocking of alpha4beta7 gut-homing integrin during acute infection leads to decreased plasma and gastrointestinal tissue viral loads in simian immunodeficiency virus-infected rhesus macaques. J Immunol. 2011;186(2):1044– 1059.

41. Hua CK, Ackerman ME. Increasing the Clinical Potential and Applications of Anti-HIV Antibodies. Front Immunol. 2017;8:1655.

42. Morris L, Mkhize NN. Prospects for passive immunity to prevent HIV infection. PLoS Med. 2017;14(11):e1002436.

43. Nishimura Y, Martin MA. Of Mice, Macaques, and Men: Broadly Neutralizing Antibody Immunotherapy for HIV-1. Cell Host Microbe. 2017;22(2):207–216.

44. Pegu A, Hessell AJ, Mascola JR, Haigwood NL. Use of broadly neutralizing antibodies for HIV-1 prevention. Immunol Rev. 2017;275(1):296–312.

45. Wagh K, Seaman MS, Zingg M, et al. Potential of conventional & bispecific broadly neutralizing antibodies for prevention of HIV-1 subtype A, C & D infections. PLoS Pathog. 2018;14(3):e1006860.

46. Barouch DH, Whitney JB, Moldt B, et al. Therapeutic efficacy of potent neutralizing HIV-1-specific monoclonal antibodies in SHIV-infected rhesus monkeys. Nature. 2013;503(7475):224–228.

47. Nishimura Y, Gautam R, Chun TW, et al. Early antibody therapy can induce long-lasting immunity to SHIV. Nature. 2017;543(7646):559–563.

48. Schoofs T, Klein F, Braunschweig M, et al. HIV-1 therapy with monoclonal antibody 3BNC117 elicits host immune responses against HIV-1. Science. 2016;352(6288):997–1001.

49. Lu CL, Murakowski DK, Bournazos S, et al. Enhanced clearance of HIV-1-infected cells by broadly neutralizing antibodies against HIV-1 in vivo. Science. 2016;352(6288):1001– 1004.

50. Animal Welfare Act and Regulation of 2001. In: Code of Federal Regulations t, chapter 1, subchapter A: animals and animal products ed. Beltsville, MD: U.S. Department of Agriculture.

51. Guerra-Perez N, Aravantinou M, Veglia F, et al. Rectal HSV-2 Infection May Increase Rectal SIV Acquisition Even in the Context of SIVDeltanef Vaccination. PLoS One. 2016;11(2):e0149491.

