## Supplementary material for "Protection of rhesus macaques against vaginal SHIV challenges by VRC01 and an anti-α_4_β_7_ antibody"

FIGURE S1

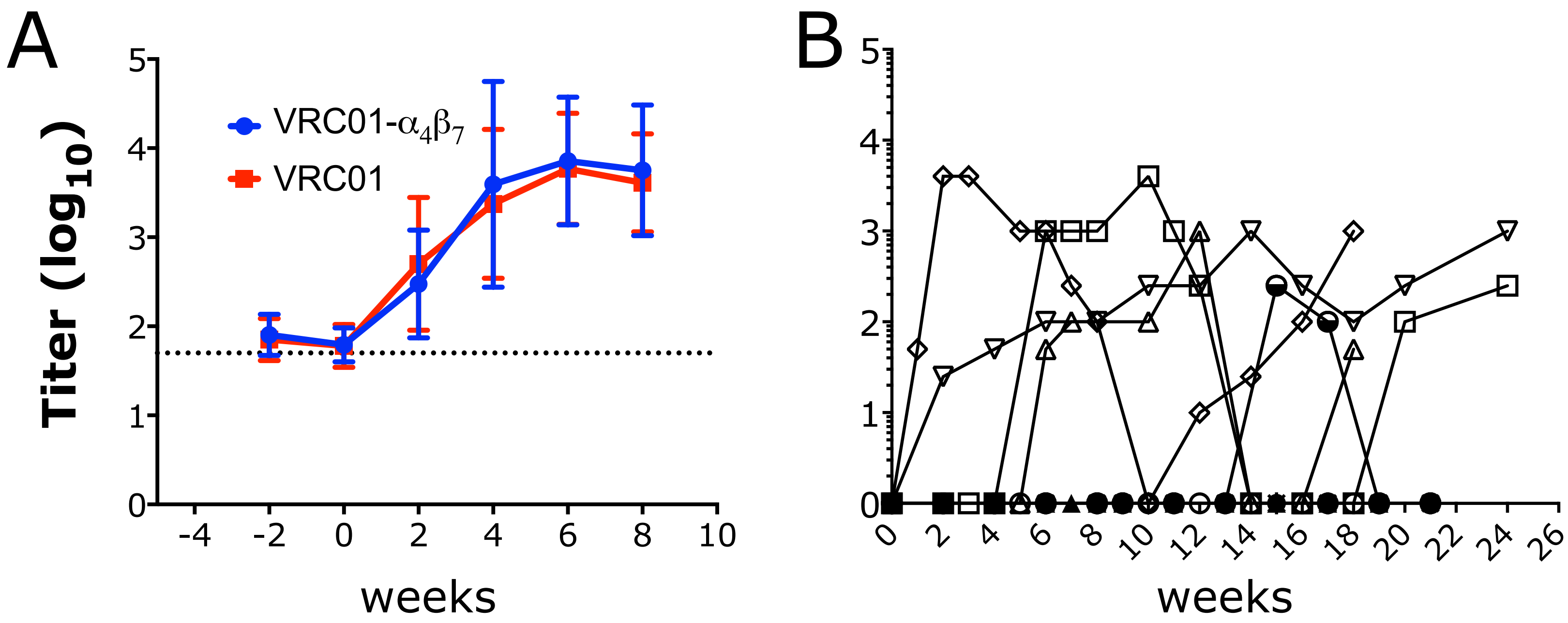


**Fig S1. Anti-drug antibodies levels.** A) Mean ± SEM of the endpoint titers of anti-VRC01 antibodies in the VRC01-alone and VRC01-$\alpha_4\beta_7$ group are shown at baseline and for the first 8 weeks past-infusion. B) Endpoint titers of anti-Rh-$\alpha_4\beta_7$ antibodies are shown for the VRC01-$\alpha_4\beta_7$ group from baseline to the necropsy.

FIGURE S2

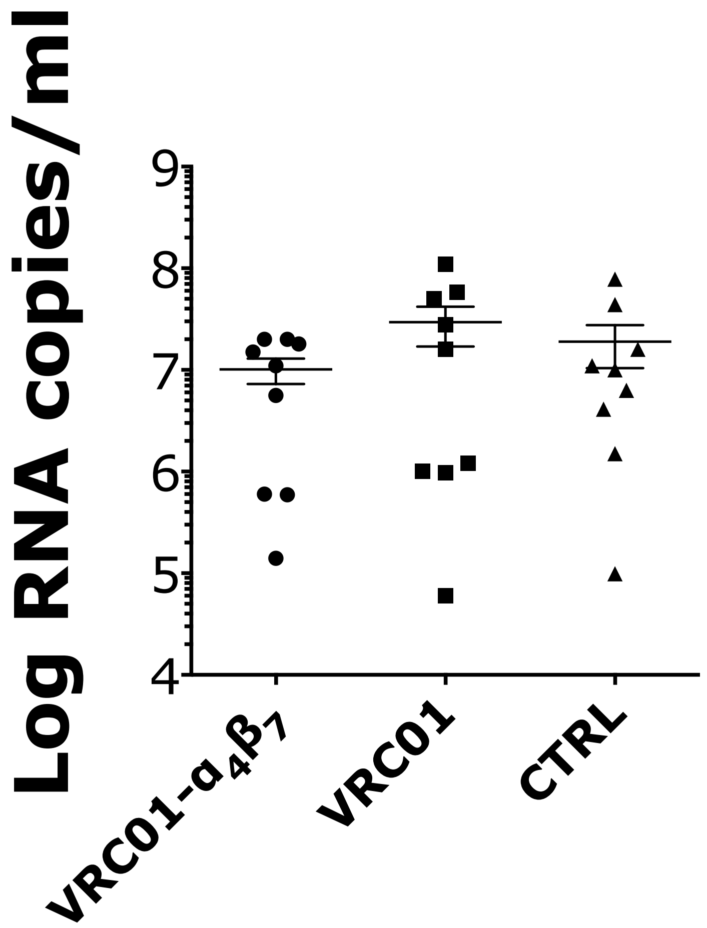


**Fig S2. No difference in peak plasma viral load among the treatment groups**. Highest level of SIV RNA copies in plasma reached within the first 5 weeks of infection in each animal is shown.

FIGURE S3

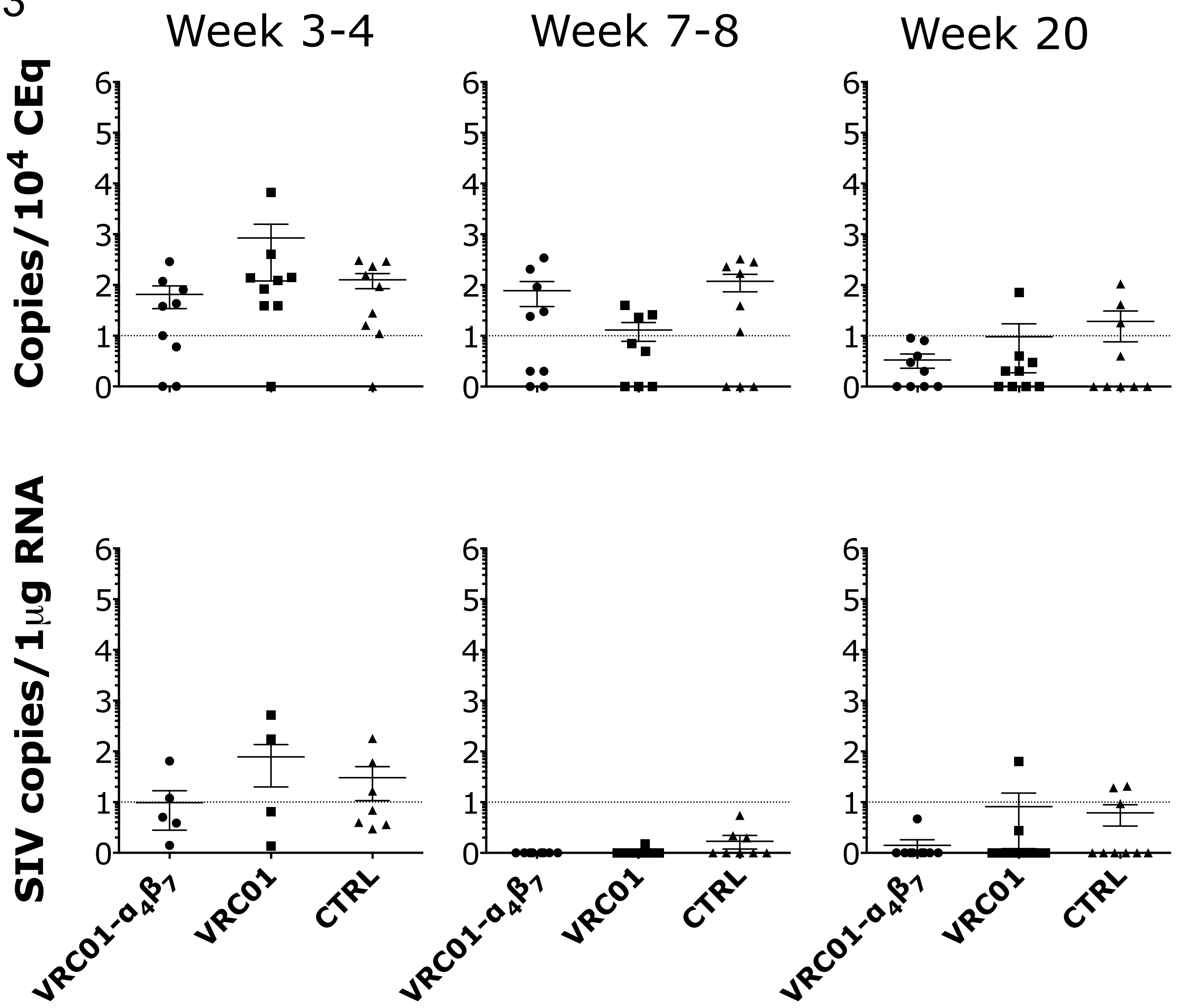


**Fig S3. No difference in vaginal tissue viral load among the treatment groups.** Copies of SIV DNA (A) and RNA (B) from vaginal biopsies at the indicated times after infection were quantified by gag-qPCR (normalized on albumin content) and by RT-qPCR (normalized on RNA content) respectively. The dotted line indicates the lower limit of detection (LLOD) of the assay.

FIGURE S4

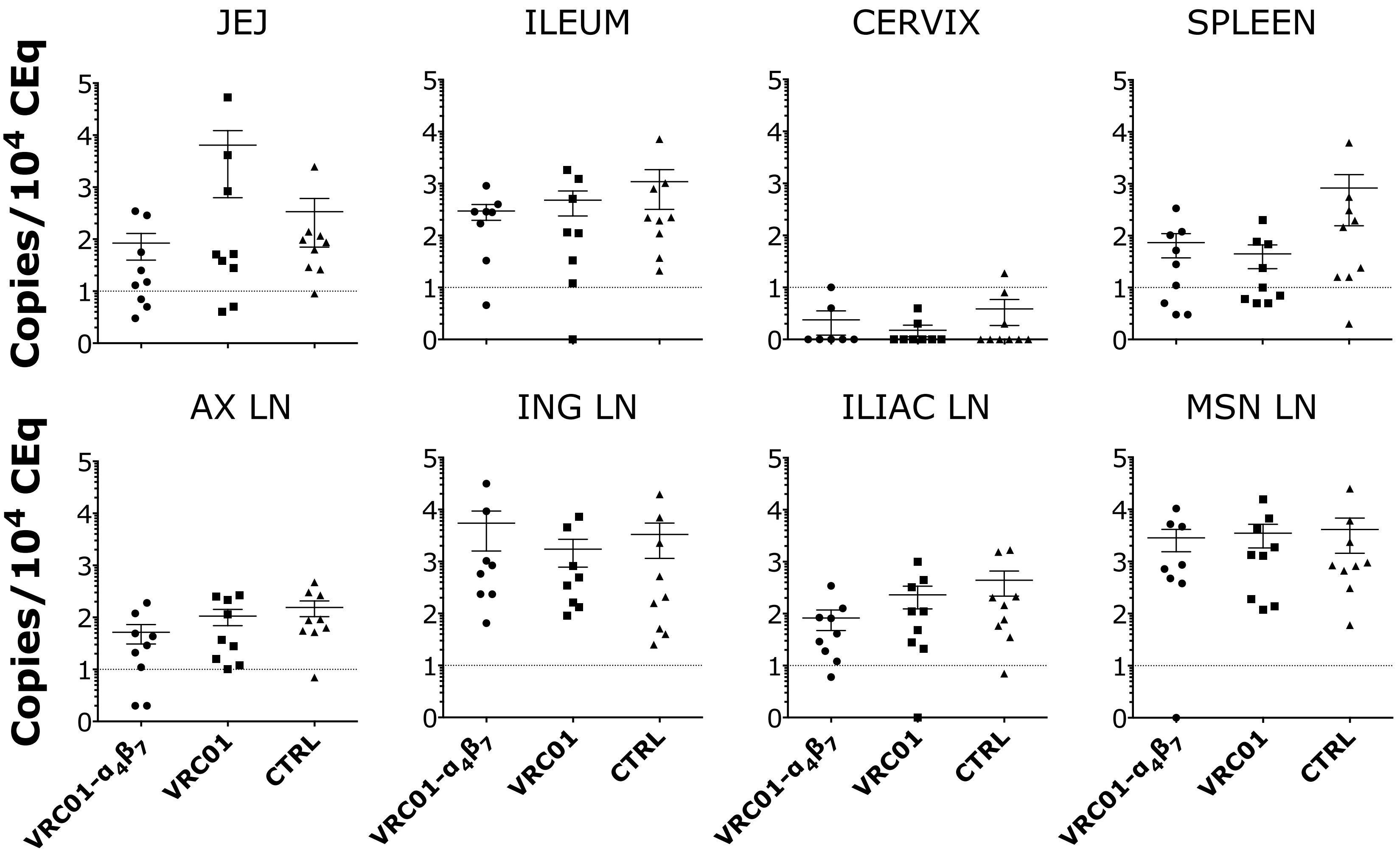


**Fig S4. SIV DNA loads in different tissues at necropsy.** Viral DNA loads in each tissue were measured by SIV gag qPCR and normalized on albumin copies. The dotted line indicates the lower limit of detection (LLOD) of the assay.

FIGURE S5

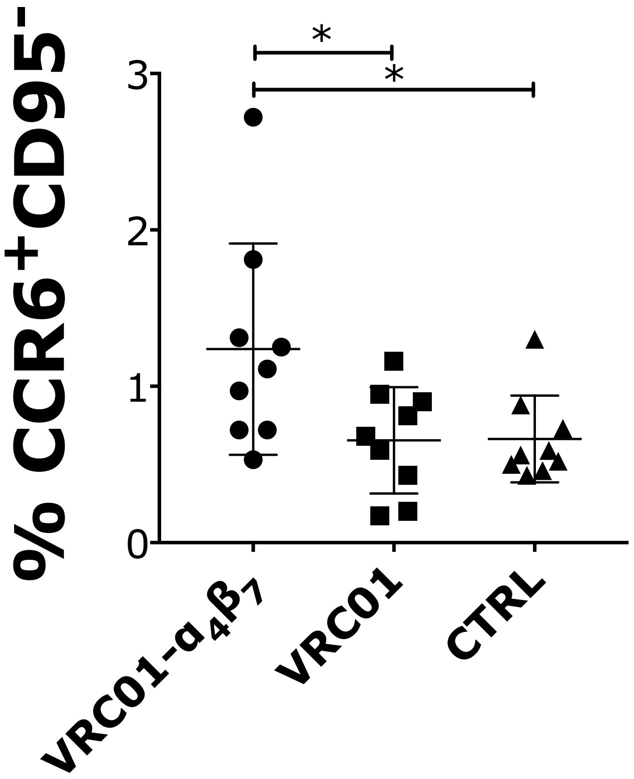


**Fig S5 Rh-$\alpha_4\beta_7$-VRC01-treated macaques have higher levels of circulating $CCR6^+$ $CD4^+$ T cells in the chronic phase.** Around week 20 p.i. blood T cells were phenotyped by flow cytometry. The frequency of the subset that significantly differed among the treatment group is shown. The results of the Dunn's multiple comparisons post-hoc test (after the Kruskal-Wallis test controlled for multiple comparisons) and the Mann-Whitney test to compare the treatment groups between each other are shown ($p$-value of * $\alpha<0.05$, was considered significant).

### FIGURE S6

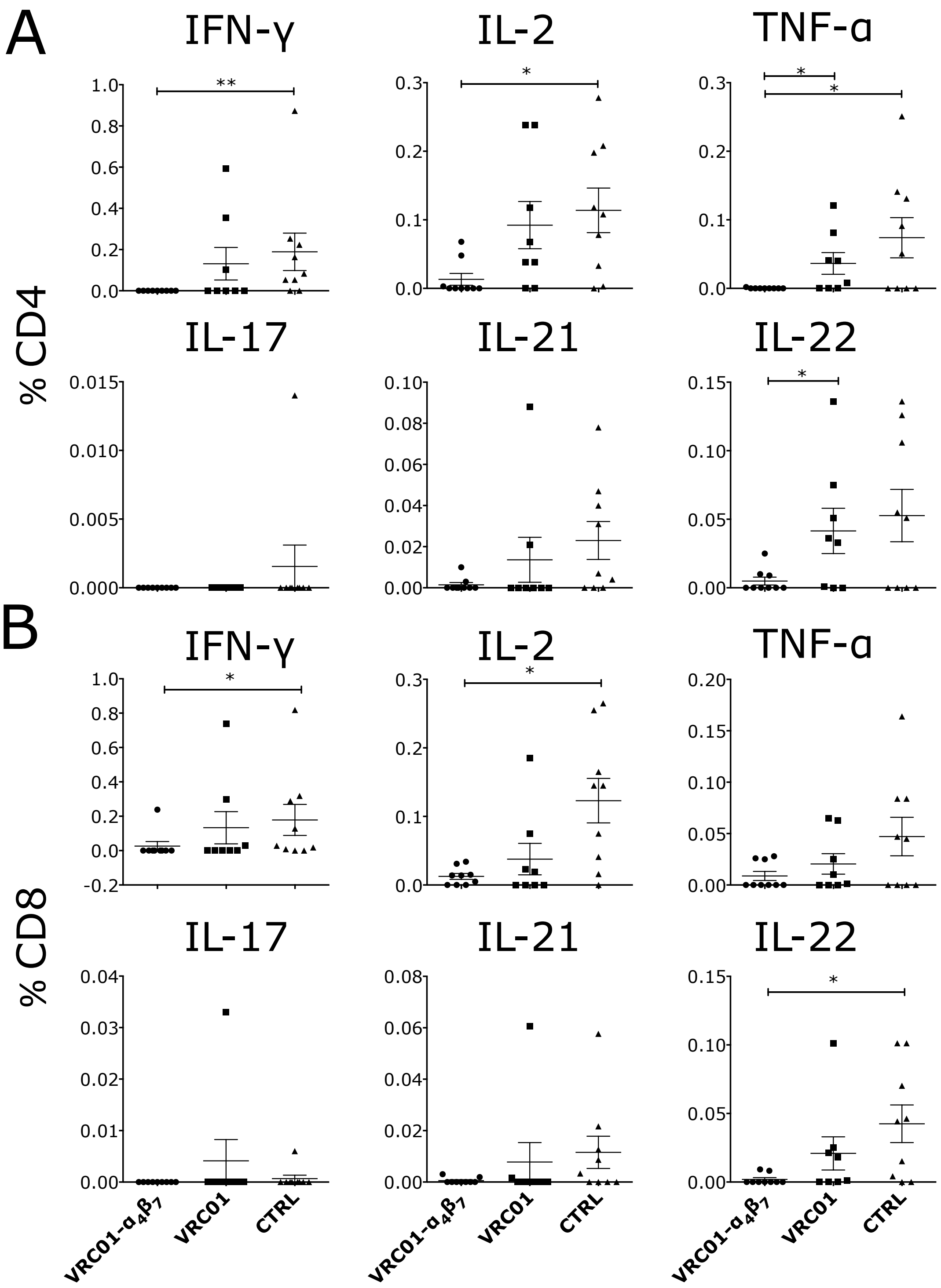


**Fig S6 Blood T cell responses against the consensus B envelope peptide pool.** PBMCs isolated around 18 weeks post infection were stimulated with pooled 15-mer peptides with an 11aa overlap from the consensus B envelope protein for 5hrs. The frequency of cells secreting the indicated cytokines are shown for the $CD4^+$ and $CD8^+$ T cell subsets after subtraction of the baseline values (in absence of peptides). The results of the Dunn's multiple comparisons post-hoc test (after the Kruskal-Wallis test controlled for multiple comparisons) and the Mann-Whitney test to compare the treatment groups between each other are shown (*p*-value of * $\alpha<0.05$, ** $\alpha<0.01$ and *** $\alpha<0.001$ were considered significant).

### FIGURE S7

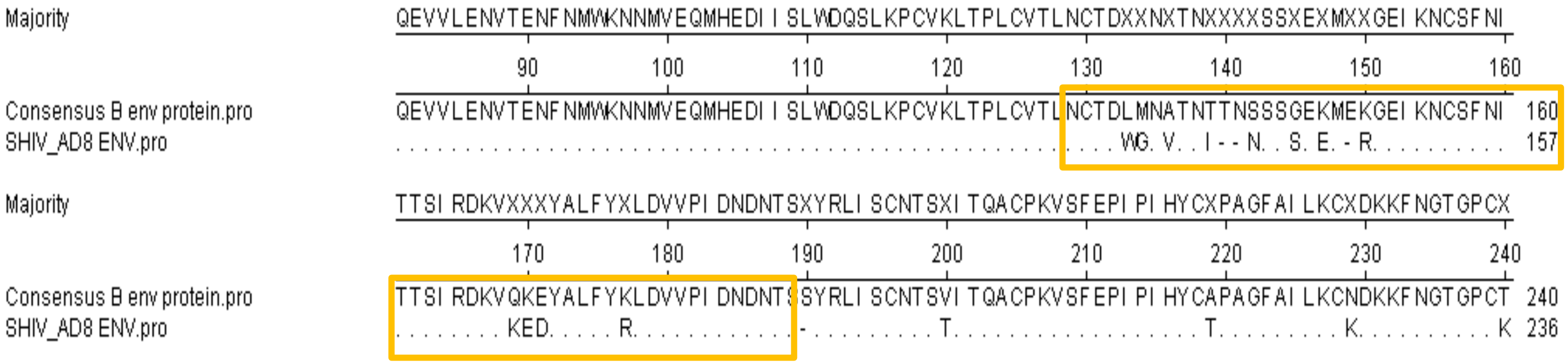


**Fig S7. Alignment of the V1-V2 region of the consensus B and SHIV$_{AD8-EO}$ envelope sequences.** Peptides of 20aa (overlapping 14aa) spanning the region shown in yellow were synthetized and used to probe T cell and antibody responses.

FIGURE S7

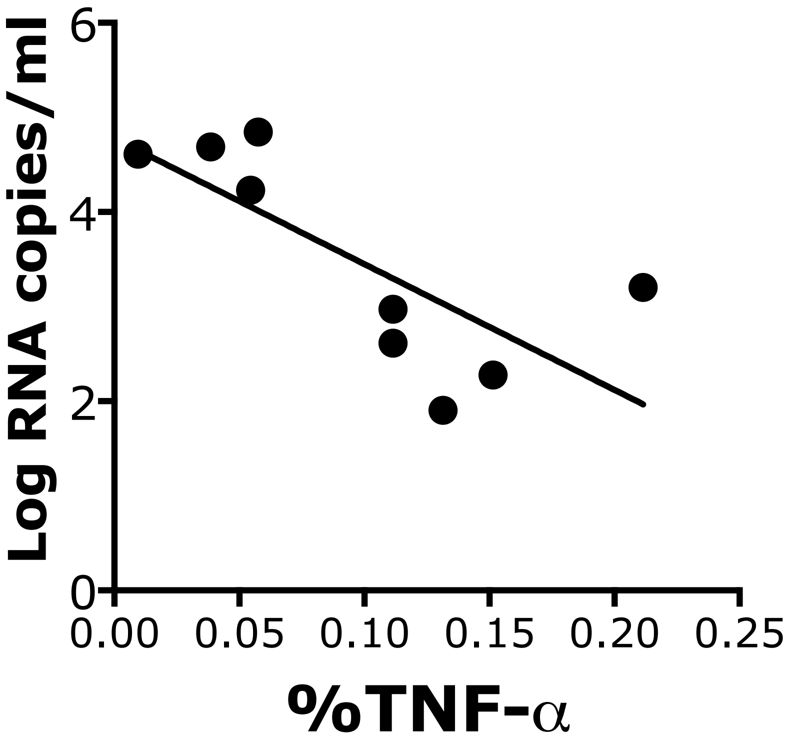


**Fig S7. CD8+ T cell responses to the V1V2 peptides correlated with viral load.** The frequencies of CD8+ T cells producing TNF-α in response to V1V2 peptides in the blood of VRC01-Rh-$\alpha_4\beta_7$ (shown in Fig 5B) inversely correlate with viral loads.

FIGURE S9

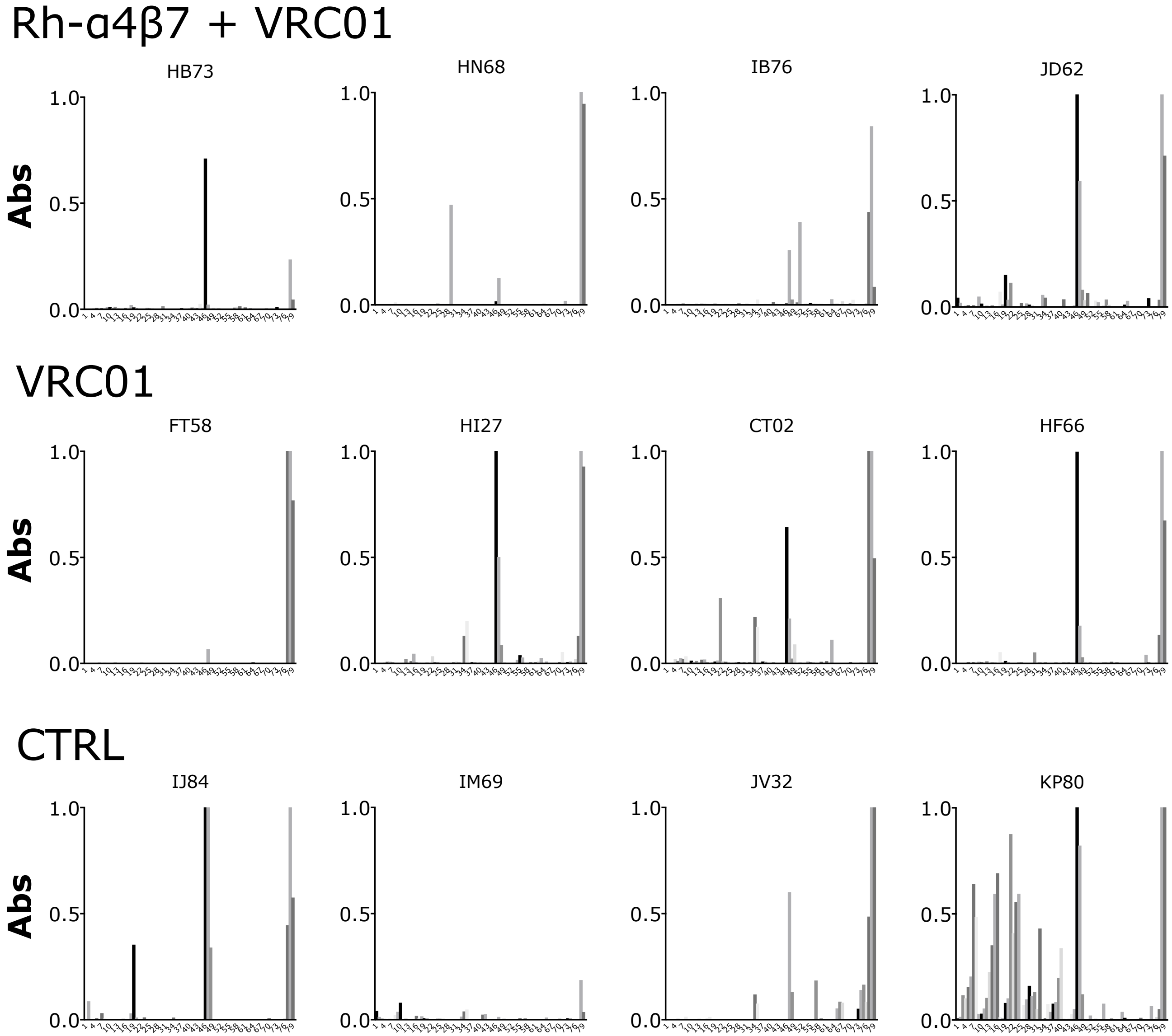


**Fig S9. Peptide scan.** Serum from 4 animals from each treatment group was analyzed by peptide scan against consensus B envelope peptides. 7 SHIV-AD8-specific peptides replaced the corresponding peptides in the V1-V2 loop region.

| PBMC acute | Color | Clone | Manufacturer |
|---|---|---|---|
| CD3 | V450 | SP34-2 | BD Biosciences |
| CD4 | BUV395 | L200 | BD Biosciences |
| CD95 | PercP-Fluor710 | DX2 | eBioscience |
| CCR6 | PE-Dazzle 594 | G034E3 | BioLegend |
| NKG2A | PE | REA110 | Miltenyi |
| alpha4 | AF700 | 7.2R | Novus Biologicals |
| CXCR3 | AF488 | G025H7 | BioLegend |
| IL-17A | PE-Cy7 | Ebio64DEC17 | eBioscience |
| P27 | AF647H | 2F12 | NIAIDS |
| CCR7 | BV605 | G043H7 | BioLegend |
| IFNg | APC-efluor780 | 4S.B3 | eBioscience |

| RECTAL acute | Color | Clone | Manufacturer |
|---|---|---|---|
| CD3 | V450 | SP34-2 | BD Biosciences |
| CD4 | BUV395 | L200 | BD Biosciences |
| NKp44 | PercP-Cy5.5 | P44 | BioLegend |
| CCR6 | PE-Dazzle 594 | G034E3 | BioLegend |
| alpha4 | PE | 7.2R | Novus Biologicals |
| CXCR3 | AF488 | G025H7 | BioLegend |
| IL-17A | PE-Cy7 | Ebio64DEC17 | eBioscience |
| NKG2A | APC | REA110 | Miltenyi |
| CD20 | AF700 | 2H7 | BD Biosciences |
| IgA | In house APC-Cy7 | 10F12 | NHPR center |

| PBMC stimulation | Color | Clone | Manufacturer |
|---|---|---|---|
| CD3 | V450 | SP34-2 | BD Biosciences |
| CD8 | PE-CF594 | RPA-T8 | BD Biosciences |
| CD4 | BUV395 | L200 | BD Biosciences |
| NKG2A | PE Vio 770 | REA110 | Miltenyi |
| IL-17A | APC-eFluor 780 | eBio64DEC17 | eBioscience |
| IFN-gamma | AF700 | B27 | BD Biosciences |
| IL-2 | Brilliant Violet 605 | MQ1-17H12 | BioLegend |
| IL-21 | APC | 3A3-N2 | BioLegend |
| IL-22 | PerCP-eFluor 710 | IL22JOP | eBioscience |
| TNF alpha | FITC | MAb11 | BioLegend |

| Tissue stimulation | Color | Clone | Manufacturer |
|---|---|---|---|
| CD3 | V450 | SP34-2 | BD Biosciences |
| CD8 | PE-CF594 | RPA-T8 | BD Biosciences |
| CD4 | BUV395 | L200 | BD Biosciences |
| NKp44(CD336) | PerCP-Cy5.5 | P44-8 | BioLegend |
| CCR6 (DcR2) | PE-Cy7 | G034E3 | BioLegend |
| TNF alpha | FITC | MAb11 | BioLegend |
| IL-17A | APC-eFluor 780 | eBio64DEC17 | eBioscience |
| IFN-gamma | Alexa Fluor 700 | B27 | BD Biosciences |
| IL-2 | BV 605 | MQ1-17H12 | BioLegend |
| IL-21 | PE | 3A3-N2 | BioLegend |
| p27 | APC | 2F12 | |

| PBMC (chronic) | Color | Clone | Manufacturer |
|---|---|---|---|
| CD3 | Alexa Fluor 700 | SP34-2 | BD Biosciences |
| CD4 | BUV395 | L200 | BD Biosciences |
| CXCR5 | FITC | 710D82.1 | NHP |
| CXCR3 | PerCP-Cy5.5 | G025H7 | BD Biosciences |
| CD95 | V450 | DX2 | BD Biosciences |
| CD127 | PE-Cy7 | eBioRDR5 | eBioscience |
| CD25 | APC-eFluor 780 | BC96 | BioLegend |
| CCR6 | PE-Dazzle 594 | G034E3 | BioLegend |
| CD103 | PE | B-Ly7 | eBioscience |
| CD69 | Brilliant Violet 605 | FN50 | BD Biosciences |
| P27 | APC | 2F12 | NIAIDS |

| LN (chronic) | Color/Format | Clone | |
|---|---|---|---|
| CD3 | Alexa Fluor 700 | SP34-2 | BD Biosciences |
| CD4 | BUV395 | L200 | BD Biosciences |
| CXCR5 | FITC | 710D82.1 | NHPR program |
| CXCR3 | PerCP-Cy5.5 | G025H7 | BD Biosciences |
| CD95 | V450 | DX2 | BD Biosciences |
| CD127 | PE-Cy7 | eBioRDR5 | eBioscience |
| CD25 | APC-eFluor 780 | BC96 | BioLegend |
| CCR6 | PE-Dazzle 594 | G034E3 | BioLegend |
| PD-1 | PE | EH12.2H7 | BioLegend |
| CD69 | Brilliant Violet 605 | FN50 | BD Biosciences |
| P27 | APC | 2F12 | NIAIDS |

**Fig S10. Panels used for flow cytometry analysis of cell subset and T cell responses.** Acute analysis were done with samples collected at week 3 or 4 post-infection, while chornic samples were collected from week 18 to 22 post-infection.
